# A proteomic analysis of the PHF-forming tau fragment (tau297-391) following uptake into differentiated human neuronal SHSY5Y cells

**DOI:** 10.1101/2025.11.03.684805

**Authors:** Alice Copsey, Sebastian S. Oakley, Mahmoud B. Maina, Robert Parker, Karen E. Marshall, Charles R. Harrington, John M.D. Storey, Claude M. Wischik, Louise C. Serpell

**Affiliations:** Sussex Neuroscience, School of Life Sciences, University of Sussex, Falmer, East Sussex, UK; Biomedical Science Research and Training Centre, Yobe State University, Damaturu, Yobe State, Nigeria; Centre for Immuno-Oncology, Nuffield Department of Medicine, University of Oxford, Oxford, OX3 7DQ, UK; Institute of Medicine, Medical Sciences and Nutrition, University of Aberdeen, Aberdeen, UK; TauRx Therapeutics Ltd., Aberdeen, UK; Department of Chemistry, University of Aberdeen, Aberdeen, UK

**Keywords:** tau, Alzheimer’s disease, tauopathies, proteome, PRDX6

## Abstract

Tau self-assembly and intracellular deposition is associated with a group of neurodegenerative diseases called tauopathies, which include Alzheimer’s disease (AD) and Pick’s disease. Here, we measured the proteome response in human neuronal cells (differentiated SH-SY5Y) following addition of a spontaneously amyloidogenic region of tau known as dGAE (tau297-391) which forms AD-like paired helical filaments in vitro and proteomic analysis showed increased endogenous tau expression. Further interactome analysis uncovered increased association between tau and proteins associated with nuclear chromatin, the nucleolus, and the spliceosome as well as the thiol-peroxidase, PRDX6 alongside an increase in reactive oxygen species (ROS). This work highlights a method to identify proteome pathways that may play an important role in the development of tau pathology.

## Introduction

Tau is a microtubule-associated protein responsible for promoting and stabilising the microtubule network. Alternative splicing of the MAPT gene located on chromosome 17 results in the expression of the six major isoforms in the central nervous system, which consist of 0, 1 or 2 N-terminal regions and three or four microtubule-binding regions (MTBR) [1] [2]. In addition to its role in the microtubule network, it has been suggested that tau is involved in functions within the nucleus. These include protection against reactive oxygen species (ROS) [3], ribosome stability and miRNA activity [4] and nucleolar transcription and nucleolar stress response [5, 6]. However, in a group of neurodegenerative diseases, termed tauopathies, tau undergoes pathological processes which results in tau assembling into intracellular tau aggregates. These tau aggregates comprise of disease-specific amyloid fibrils resulting from pathological tau self-assembly [1] and their accumulation is closely associated with the neurodegeneration and the cognitive decline observed in people living with these diseases. Oxidative stress contributes to the pathology of neurodegenerative diseases, including tau pathology in tauopathies [7]. Elevated OS may be an early trigger in tau pathogenesis preceding tau aggregation or be a consequence of tau misfolding and aggregation [8]. It has been speculated that OS generated at early stages of AD induces the formation of pathological tau, which promotes mitochondrial impairment resulting in increased OS generation leading to neuronal death [8].

The processes whereby physiological tau is modified into a form of tau that promotes self-assembly and early stages of pathology remain uncertain. Post-translational modifications (PTMs), such as phosphorylation and truncation, are thought to play a crucial role in both the physiological function of tau and its pathological assembly [9]. An *in vitro* model has shown that tau297-391 (termed dGAE) spontaneously self-assembles into amyloid fibrils that exhibit the same macromolecular structure to paired helical filaments (PHFs) in AD and type II filaments in CTE [10] [11]. This fragment of tau was first identified as a species of tau isolated from proteolytically stable core preparations of PHFs from AD brains [12]. Soluble dGAE can accumulate within intracellular compartments of differentiated human neuroblastoma cells and induce changes in the distribution and species of tau within the cells [13].

In this study, we have investigated the downstream effects of soluble dGAE on the proteome of differentiated human neuroblastoma SH-SY5Y (dSH). We have utilised mass spectrometry (MS) analysis to examine changes in the proteome and tau interactome in response to a 24-h incubation of soluble dGAE. The proteomic analysis identified an increase in relative abundance of proteins associated with AD pathology, including tau protein itself. Further analysis of the tau interactome after treatment with dGAE showed an enrichment in tau interactors associated with nuclear chromatin, the nucleolus, and the spliceosome. There was also a small but significant increase in ROS after 24 hours incubation with dGAE. The findings support the use of dGAE as a model to initiate tau pathology and suggest that the observed increase in tau binding to chromatin is likely the result of an increase in OS.

## Methods

### Protein expression and purification

dGAE was produced using a protocol previously described [10].

### Cell Culture

SH-SY5Y (SH) cells were cultured in Advanced DMEM /F12 (Dulbecco’s modified Eagle medium; Gibco) supplemented with 10% (v/v) foetal calf serum (FCS (Labtech, U.K.)), 2mM L-glutamine and 1% PenStrep (Gibco) in a humidified atmosphere containing 5% CO_2_. Once cells reached required confluency, the Advanced DMEM /F12 media was removed and refreshed with Advanced DMEM /F12 supplemented with 1% FCS and 10µM retinoic acid and incubated in the dark in a humidified atmosphere containing 5% CO_2_ for 2 to 3 days. The retinoic acid-containing media was refreshed and cells incubated for another 2 to 3 days. Cells were washed in serum-free media to remove traces of FCS and replaced with Advanced DMEM /F12 containing 50ng/ml BDNF and incubated for a further 2 days. SH cells were fully differentiated SH (dSH) cells after the 2d BDNF treatment and were used for experimentation.

### Total protein extracts

Cells were washed with phosphate-buffered saline (PBS), scraped into PBS, pelleted by centrifugation in a cooled microcentrifuge at 10⍰000×***g*** at 4°C and resuspended in either a denaturing lysis buffer [25mM Tris-HCl (pH7.5), 4% SDS] or a non-denaturing lysis buffer [25mM TrisHCl (pH 7.5), 150mM NaCl, 1mM DTT, 1mM EDTA, 1% NP40, 0.5% sodium deoxycholate, 0.05% SDS, 1 unit Benzonase, 2mM benzamidine, 5µg/ml leupeptin, 10mM chymostatin, 1X PhosSTOP^TM^ (Roche) and 1× EDTA-free protease inhibitor cocktail (Roche)]. After lysis, cell debris was removed by centrifugation in a cooled microcentrifuge at 10⍰000×***g*** at 4°C. Protein concentration was determined using the Pierce BCA protein assay as described by the manufacturers.

### Protein digestion with trypsin to produce tryptic fragments for LC-MS analysis

On-column digests were performed using S-Trap™ micro-spin columns according to the manufacturer’s instructions. In brief, protein was bound to the column in 5% SDS lysis buffer supplemented with 50mM Tetraethylammonium bromide (TEAB). Disulphide bonds were reduced with 5mM tris(2-carboxyethyl)phosphine) (TCEP) and alkylated with 20mM *S*-Methyl methanethiosulfonate (MMTS). After acidification protein was washed with binding buffer and digested with trypsin (Promega). Peptides were eluted into 50% acetonitrile in water, dried in a speed vac and resuspended in 0.1% trifluoracetic acid in water for MS analysis.

### Liquid chromatography - mass spectrometric (LC-MS) acquisition

Samples were analyzed on either a U3000 HLPC coupled to a Q-Exactive (HFX) Orbitrap mass spectrometer (Thermo Scientific) or a Nano-elute HPLC coupled to a TimsTOF SCP mass spectrometer – (Bruker Daltonics).

For HFX, peptides were initially trapped and then separation was conducted using a 180 min acetonitrile gradient 2-35% applied over a 75μm × 50 cm PepMap RSLC C18 EasySpray column (Thermo Scientific) at a flow rate of 250nl/min. Peptides were analysed by data-dependent acquisition full-MS1 scan (60,000 resolution, 45 ms accumulation time, AGC 3×10^6^) and 10 data-dependent MS2 scans (15,000 resolution, 54 ms accumulation time, AGC 2×10^5^), with an isolation width of 1.3 m/z and normalized HCD energy of 28%, only 2-4 charge states were fragmented.

For TimsTOF SCP – (Bruker Daltonics), peptides were initially trapped and then separation was done using a 60 min gradient of 2–40% acetonitrile in 0.1% Formic acid at a flow rate of 150nL/min with an Aurora 25 cm × 75μm, 1. μm, C18 column (IonOpticks, Australia). A CaptiveSpray source (Bruker Daltonics) set to 180°C, 1500 V capillary voltage and 3 L/min dry gas was used to ionise peptides. The instrument was operated in DDA-PASEF mode with one MS survey TIMS-MS and 10 PASEF MS/MS scans acquired at a ramp time of 166ms within an ion mobility range of 1/K0 = 1.45 Vs cm2 to 0.64 Vs cm-2 and *m*/*z* range of 100–1,700. Multiply charged ions were included with a minimum threshold of 500 arbitrary units (a.u.), fragmented and re-sequenced until reaching 20,000 a.u. Collision energies were used as follows: 50 eV at 1/K0 = 1.6 Vs cm^−2^ and 20 eV at 1/K0 = 0.6 Vs cm^−2^

### Quantitative proteomics data analysis

Label free quantitative proteomics data analysis was performed in Maxquant v 1.6.1.0, utilizing human Swiss-prot database containing approximately 20,000 entries downloaded on 24/05/2018. Trypsin/P was set for the enzyme specificity with up to 2 missed cleavages allowed. Variable modifications were set to N-terminal (M+ 42.01) acetylation and oxidation (M, +15.99 Da), with a fixed modification for carbamidomethylation (C, +57.02) also set and only peptides and proteins passing 1% FDR were reported. LFQ values were used for proteomics experiments and intensity values for interactomics, and the protein groups file imported into Perseus v. for statistical analysis. For proteomics values were log 2 transformed and reverse hits were removed, we filtered based on consistency with 2/3 real values required in at least one condition for a protein to be valid. Next, normalised using width adjustment and imputed missing values from the normal distribution for each sample separately using down shift of 1.8 and a width of 0.3. A Student’s t test was used to calculate the significance and fold-changes for each protein between experimental conditions. For interactomics, data analysis was identical except data was filtered based on a data consistency of 2/2 in at least one condition. Volcano plots were plotted in Excel.

### Quantitative Western blotting

Protein extracts were separated by SDS–PAGE, transferred to 0.2µm nitrocellulose membranes and visualised with the appropriate antibodies followed by HRP-conjugated secondary antibodies (DAKO, 1:⍰2000). HRP activity was detected using the Bio-Rad Clarity^TM^ Western ECL Substrate followed by exposure using the Odyssey®Fc imaging system. Protein bands of interest were quantified using the densitometry analysis software tool.

### Immunoprecipitation

Aliquots of extract (non-denaturing) containing 500 μg of protein were incubated either with 5µg of Total tau (Sigma SAB4501821, H2A (Abcam 177308) SF3A3 (Proteintech 12070-1-AP) or PRDX6 (Proteintech 13585) antibody overnight at 4°C with the addition of 50 µl of 50% Protein A-Sepharose for the last 30 min. Recovered proteins were washed three times with IP lysis buffer (Pierce 87787) supplemented with protease and phosphatase inhibitors. After washing, bound proteins were recovered with 7M urea and 2M thiourea in 25 mM Tris–HCl (pH 7.5).

### Differential salt fractionation

Nuclear pellets were prepared using the Nuclei isolation kit (Sigma NUC101) following the procedure for attached cells. Supernatant and insoluble nuclei were retained for analysis. Differential salt fractionation of nuclear DNA was performed as outlined by C. Herrmann *et. al*. [14]. In brief, nuclei were resuspended in Mnase digestion buffer (10nM Tris pH 7.4, 2mM MgCl_2_, 0.1mM PMSF) with 1 unit of Mnase and incubated for 30 minutes at 37°C in a water bath. The digestion was stopped by the addition of 0.1 M EGTA. After centrifugation for 10 minutes at 400g, nuclei were washed in the digestion buffer supplemented with 5mM CaCl_2_. Nuclei were then fractionated using buffers with increasing salt concentration (10mM Tris pH 7.4, 2mM MgCl_2_, 2mM EGTA, 0.1% Triton X-100, NaCl 150mM-600mM). Incubations were for 30 minutes at each salt concentration, rotating at 4°C, followed by centrifugation at 400g for 10 minutes. The final pellet was suspended in RIPA buffer to release protein from actively transcribed regions of the chromatin. Eluted fractions were analysed using western blotting with antibodies specific to the nuclear fraction (H2A; Abcam 177308), supernatant (GAPDH) and Tau protein (Sigma SAB4501821).

### Immunofluorescence

SH cells (40,000) were plated on 13-mm coverslips and differentiated with RA/BDNF protocol described above. dSH cells were treated with 10µM soluble dGAE or dGAE-C322A, or equal volume of phosphate buffer as the control, in 0% FCS media for 24h before being fixed with 4% (v/v) paraformaldehyde/PBS, permeabilised with 0.1% (v/v) TritonX-100 and treated with 50mM glycine to block aldehydes. Immunostaining was done with anti-PRDX6 or anti-tau(MAPT) (Sigma-Aldrich) antibody for 1h at RT before being mounted with Prolong™ Gold Antifade Mountant with DAPI.

### Confocal microscopy

All confocal microscopy imaging was carried using a Leica SP8 confocal microscope. The instrument setting used PMT 3 and PMT Trans channels/lasers and images were acquired with a HC PLAPoCs2 63 x/1.40 oil-immersion objective lens. Samples were scanned sequentially to prevent spectral bleed through. All images were collected as Z-stacks for all channels using a step size of 0.5μm. Five to ten Z-stacks were taken for each sample.

To analyse the PRDX6 response after treatment with varying proteins, images were Z-projected to standard deviation and the punctate PRDX6 fluorescence was isolated using ‘intermodes’ thresholding. A mask was created of the threshold image and watershed separation was applied to separate touching objects. Particle analysis was then performed to quantify the number of individual particles there were in the mask and the area that they covered. All results were then normalised to the number of cells within the imaged area, using the DAPI channel to manually count the number of cells. To measure changes in colocalization between endogenous tau and PRDX6 after 24h of treatment with 10µM dGAE, the BIOP JACoP plugin was utilised. This was used over other colocalization plugins because it was able to analyse each Z-slice of the stack and had thresholding settings that worked well with the images. ‘Intermodes’ thresholding was used for the PRDX6 channel images, and the ‘default’ thresholding was used for the tau channel (3R tau). The Mander’s correlation coefficient was taken for the ratio of 3R tau colocalising with PRDX6 and given as the M1 value from the readout. BIOP JACoP was also used to measure the colocalization of dGAE (E2E8 antibody) with PRDX6 using ‘intermodes’ thresholding for both channels.

### CellROX™ Green Reagent Assay

SH cells (10,000) were plated into black, clear bottom 96/well plate and differentiated with RA/BDNF protocol described above. dSH cells were treated with 10µM of soluble dGAE or equal volume of phosphate buffer as the control for 2h or 24h in 0% FCS media. 45min before the end of the incubation period, cells were treated with 5µM CellROX™ Green Reagent (Invitrogen) in 0% FCS and an additional soluble 10µM dGAE. The CellROX™ Green Reagent is a fluorogenic probe used for measuring oxidative stress. The dye is weakly fluorescent in its reduced state but exhibits a bright green fluorescence upon oxidation by ROS and subsequent binding to DNA, with absorption/emission of 485/520nm. Treatment of 7µM staurosporine for 1h was used as a positive control. Cells were then fixed with 4% (v/v) PFA before being imaged using Operetta CLS high-content analysis system (PerkinElmer, Beaconsfield, UK). Images were acquired using DAPI and FITC filters to capture the full fluorescence intensity across z-stacks, ensuring comprehensive volumetric analysis. The Harmony software’s automated analysis algorithm was utilized for precise segmentation, distinguishing nuclear from cytoplasmic signals. This allowed for quantification of the CellROX signal specifically within the nuclei, where the probe binds to DNA. Each experimental condition was independently replicated three times, analyzing a minimum of 3000 cells per replicate to ensure statistical robustness.

## Results

### Internalisation of dGAE by dSH cells results in an increase in endogenous tau protein

We have previously shown that soluble dGAE is internalised into dSH cells within 24h post-treatment [13]. To investigate the effects dGAE internalisation has on the proteome, dSH cells were incubated with 10 μM soluble dGAE for 24h and quantitative liquid chromatography mass-spectrometry (LC-MS) was used to investigate the relative abundance of proteins compared to untreated control cells. Proteomic analysis from three repeats showed that out of a total of 4447 proteins identified, only four were significantly increased post-treatment with dGAE while three were significantly decreased (>2-fold change, P=<0.05) (Fig. 1a)

**Figure 1.**
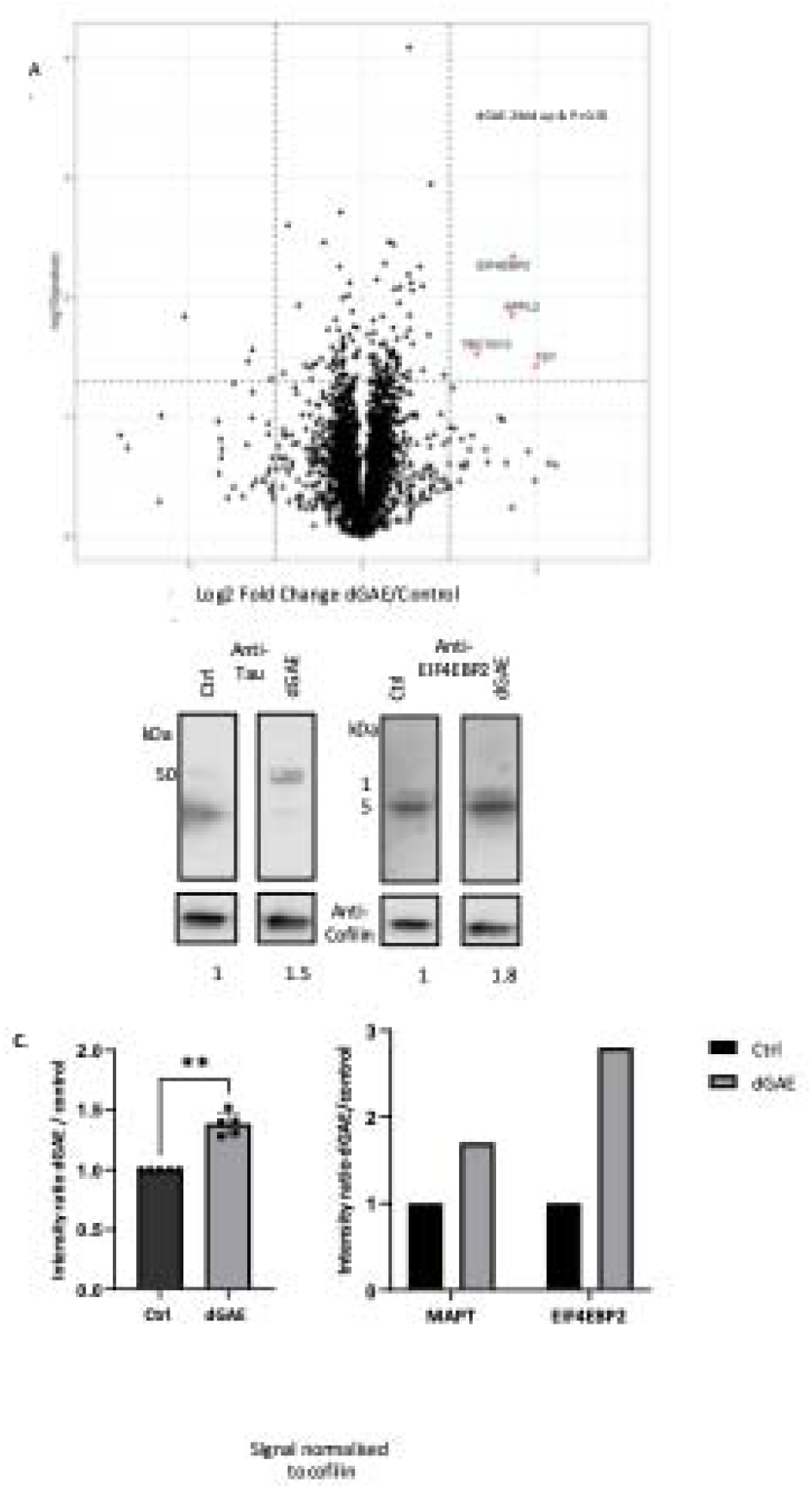
Proteomic analysis of the downstream effects of Tau after treatment of dSH cells with soluble dGAE. **A) Volcano plot showing the fold-change and P value calculated using a t test comparing protein abundances for dGAE treated and untreated cells**. Protein lysates were prepared from dSH cells 24 hours post incubation with 10µM dGAE, alongside untreated control cells. The cells were lysed using 4% SDS in 50mM Tris pH8 plus Benzonase and prepared for LC-MS analysis using S Trap™ spin columns. Protein was reduced, alkylated, and digested with trypsin to produce tryptic peptides for analysis by LC-MS as described in the material and methods. Raw mass spectra were identified and quantified using Maxquant at 1% FDR. The graph shows the gene name for proteins that were at least 2-fold enriched (Log 2 of -1, +1) with a P value of <0.05 (-Log10 P value of 1.3). **B) Western blot analysis of protein lysates prior to digestion**. Aliquots of protein prior to digestion were resolved by SDS-PAGE and visualised by immunoblotting using antibodies to tau (Sigma SAB4501831), EIF4EBP2 (cell signalling 2845) and cofilin (Sigma C8736). Bar chart shows fold-change of tau and EIF4EBP2 after treatment with dGAE compared to untreated control cells. The protein signal was normalised to the loading control cofilin. **C) Bar plot showing quantification of western blot analysis (n=4)**. Levels of Tau expression in SH-SY5Y cells post-treatment with dGAE were quantified relative to cofilin and plotted using GraphPad Prism.

The mitochondrial essential enzyme thiosulphate cyanide sulfurtransferase (TST) was increased 4-fold. This protein has been linked to oxidative stress in neurons with possible links to AD [15]. The endosomal trafficking protein APPL2, which also responds to oxidative stress [16], was increased 3.1-fold. TBC1D13, a GTPase activating protein for Rab35, was increased 3.2-fold. Rab proteins are key regulators of intracellular trafficking many of which have been implicated in AD [17]. The translation repressor EIF4EBP2, which acts as a regulator of synapse activity, learning and memory formation [18] was increased 2.8-fold. This gene has also been associated with AD [19].

Proteins that were decreased after treatment with dGAE relative to untreated control included TMED5, which is involved in Golgi ribbon formation; LAMTOR2, which is associated with the cytoplasmic fate of late endosomes and lysosomes; and MIPEP, which is involved in processing specific classes of nuclear-encoded proteins which are targeted to the mitochondrial matrix or inner membrane.

Interestingly, the proteomics data additionally detected a 0.4-fold increase in the abundance of tau protein in cells after incubation with dGAE (P= 0.001) (Fig. 1a). Using total cell extracts from this experiment, the increase in endogenous tau and another target, EIF4EBP2, was confirmed using quantitative western blotting (Fig. 1b). The increase in tau was further confirmed in subsequent western blotting experiments which consistently showed an increase in tau expression 24 hours post-treatment of dSH cells with 10µM dGAE (Fig. 1c, Figure S1).

### Incubation of dSH cells with dGAE resulted in an increase in tau binding to nucleolar, spliceosome and chromatin proteins

To further investigate cellular events relating to tau after dGAE treatment, changes to the tau interactome were explored using co-immunoprecipitation combined with LC-MS. Tau immuno-precipitates were prepared from dGAE treated and untreated dSH cells (Figure S2). Quantitative LC-MS was used to identify proteins whose abundance was altered in immuno-precipitates prepared after dGAE treatment compared to untreated control cells. Proteins that were increased or decreased two-fold or more after treatment with dGAE (P value of <0.05) were selected. From a total of 951 proteins, 39 were significantly increased after internalisation of the peptide (Fig 2A, Table S1). A protein-protein interaction (PPI) network of the enriched proteins was generated using the Search Tool for the Retrieval of Interacting Genes/Proteins (STRING) database [20]. Functional enrichments for the uploaded proteins are shown for Gene Ontology terms, Chromosome (GO:0005694), Nucleolus (GO:0005730), and Spliceasomal complex (GO:0005681) (Fig. 2b), with additional enrichments shown in the Supplementary (Table S2). Proteins whose abundance was reduced relative to wild type are listed in the Supplementary (Table S3).

**Figure 2.**
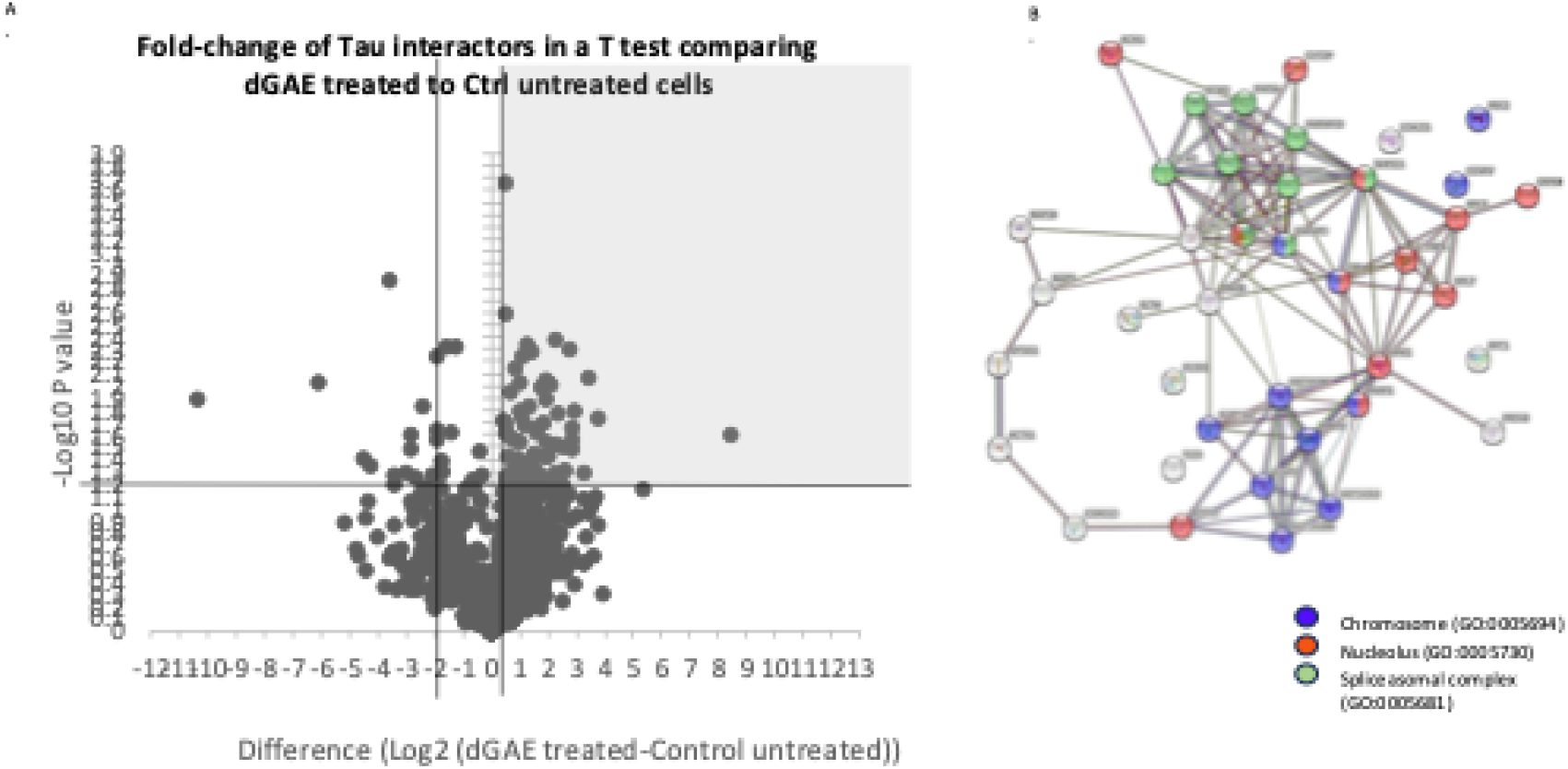
Identification of tau interactors that were enriched after treatment of dSH cells with soluble dGAE. **A) Volcano plot to show the fold-change and P value calculated using a t test comparing tau interactors plus and minus treatment with dGAE**. Protein lysates were prepared for immunoprecipitation from differentiated SH-SY5Y cells 24 hours post-incubation with 10µM dGAE, alongside untreated control cells. Cells were lysed in IP buffer and incubated with total tau antibody (Sigma SAB4501831) as described in the materials and methods. Precipitates were eluted into 8M Urea, 2M Thio-Urea and prepared for LC-MS analysis using S Trap™ spin columns. Protein was reduced, alkylated, and digested with trypsin to produce tryptic peptides for analysis by LC-MS as described in the material and methods. Raw mass spectra were identified and quantified using Maxquant 1% FDR. The highlighted area of the graph shows the proteins that were at least two-fold higher in the dGAE treated samples with a p value of <0.05. **B) Protein-Protein Interaction (PPI) network of proteins enriched at least two-fold in the dGAE treated samples with a p value of <0.05**. The data was generated using STRING [20], reporting a network showing connections for the uploaded proteins. Shows GO terms for selected enriched categories with colour code.

The interaction between tau and proteins identified as part of the tau interactome was confirmed by reverse co-immunoprecipitation. The nucleolar protein Parp1, the spliceasomal protein SF3A3 and the chromatin protein H2A were selected for validation to represent the three GO categories of interest (Table S2). The interaction between tau and selected proteins was confirmed in dSH cells 24 hours post-treatment with dGAE using immunoprecipitation of the target protein followed by western blotting (Figure S3). Tau was present in Parp1 and SF3A3 immuno-precipitates, but the interaction between tau and H2A was not confirmed by western blotting. This may have been due to the sensitivity of the antibody used.

### Tau binds to chromatin in dSH cells after internalisation of dGAE

The quantitative MS data showed an increase in tau binding to proteins associated with the chromatin after internalisation of dGAE (Fig. 2b, 3a). Interaction between tau and histone proteins was confirmed using H2A co-precipitation followed by LC-MS. H2A immuno-precipitates were prepared from dGAE treated and untreated dSH cells from three experiments as described in the materials and methods. Quantitative LC-MS was used to detect changes in tau protein co-precipitating with H2A. Three H2A proteins of interest were selected from the resulting MaxQuant dataset. MAPT was significantly increased relative to all three of the H2A proteins after treatment with dGAE compared to untreated controls, confirming an increased association between tau and H2A histone proteins after internalisation of dGAE (Fig. 3b).

**Figure 3.**
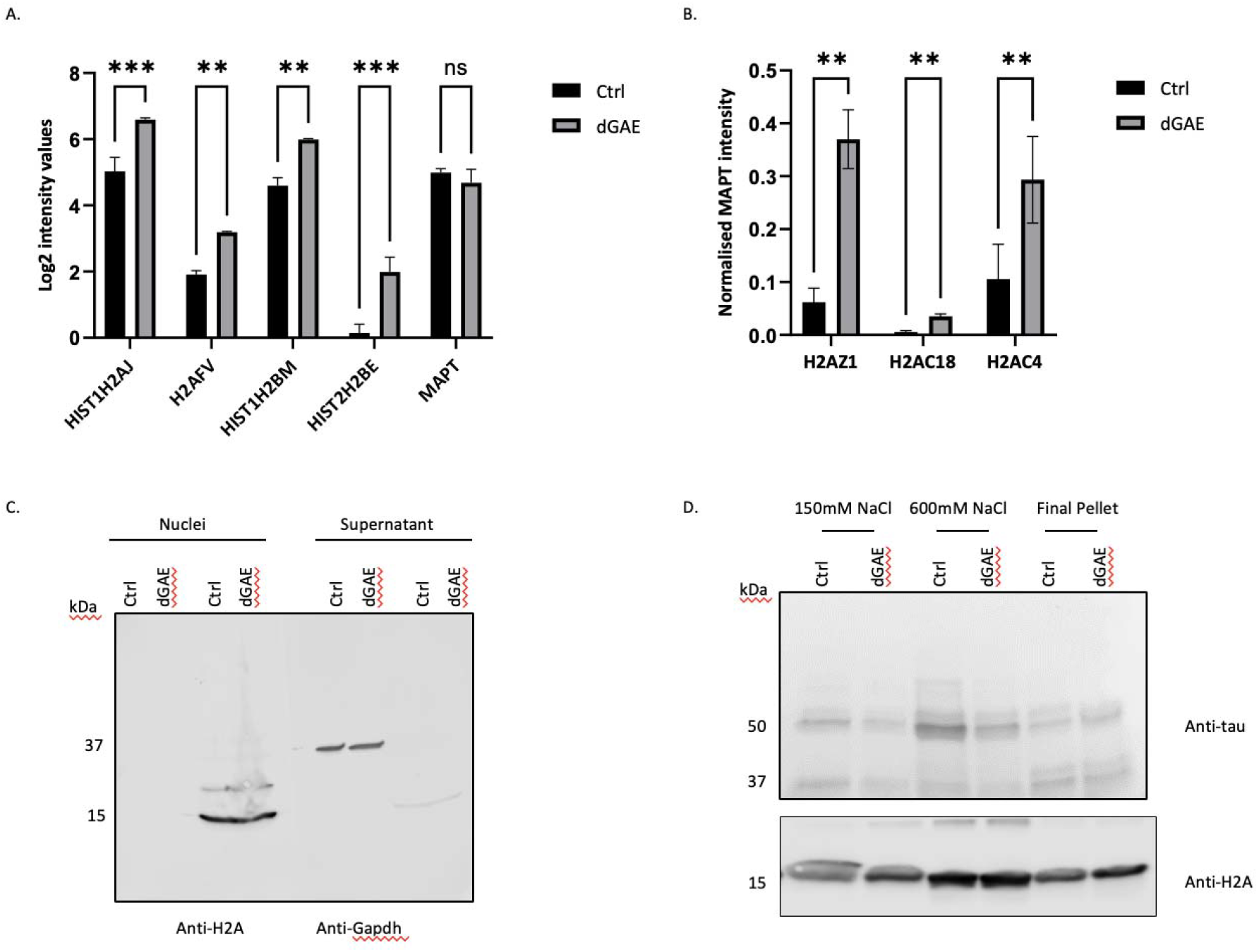
Histone proteins enriched in tau immuno-precipitates after treatment of dSH cells with soluble dGAE. A) Intensity values for histone proteins enriched in tau immuno-precipitates. Histone proteins, which were identified as enriched in tau immuno-precipitates post-treatment with dGAE (Fig. 2), were selected from the data and intensity values were plotted using GraphPad Prism alongside intensity values for MAPT (Tau protein). B) Tau co-purified with H2A immuno-precipitates. Protein lysates were prepared for immunoprecipitation from dSH cells 24 hours post incubation with 10µM dGAE, alongside untreated control cells. Cells were lysed in IP buffer and incubated with H2A antibody (Abcam 177308) as described in the materials and methods. Precipitates were eluted into 8M urea, 2M thiourea and prepared for LC-MS analysis using S Trap™ spin columns. Protein was reduced, alkylated, and digested with trypsin to produce tryptic peptides for analysis by LC-MS as described in the material and methods. Raw mass spectra were identified and quantified using Maxquant 1% FDR. The H2A proteins were selected using a cross-reactivity score threshold of 85% and greater for sequence homology between the H2A antibody and the H2A proteins in the dataset. Intensity values for MAPT in the individual immuno-precipitates were normalised to the intensity value of each H2A protein in the same sample. The normalised MAPT intensity was plotted for control and treated samples using GraphPad Prism. C) Nuclei were purified from dSH whole cell lysates using the Nuclei EZ prep kit (Merck) according to the manufacturer’s instructions for adherence cells. Nuclei and supernatant fractions were confirmed by western blotting using antibodies to marker proteins H2A (nuclei) and GAPDH (supernatant). D) Tau protein associated with both euchromatin and heterochromatin in addition to actively transcribed regions of the chromosome. Differential salt fractionation of nuclei from dSH cells was used to detect chromatin-associated tau protein. Chromatin was salt fractionated as described in the material and methods. Proteins from the 150mM salt fraction (euchromatin), the 600mM salt fraction (heterochromatin) and the final pellet (actively transcribed regions) were separated by SDS-PAGE and visualised by immunoblotting using antibodies to tau (Sigma SAB4501831) and H2A (Abcam 177308).

Differential salt fractionation of nuclei prepared from dSH cells was used to detect chromatin-associated tau protein and to identify any changes after dGAE treatment (representative images shown in Fig. 3c, d). Tau was associated with euchromatin, heterochromatin and actively transcribed regions of the chromosome in both treated and control untreated nuclei. This further confirms that tau interacts with chromatin in the nuclei of dSH cells.

### Interaction between tau and the antioxidant PRDX6 is increased in dSH cells after internalisation of dGAE

Oxidative stress (OS) resulting from increased ROS has been linked to neurodegeneration in AD [21]. In this study, an increased level of the antioxidant PRDX6 in the tau interactome after internalisation of dGAE was observed (Fig. 2b, Table S2) and this is of particular interest as it is a strong indicator of an OS response. The interaction between tau and PRDX6 was confirmed by reverse co-immunoprecipitation (Figure S3).

Although there was an increase in PRDX6 in the tau interactome (Fig. 2b, Table S2), the relative abundance of PRDX6 was unchanged in the proteomic analysis (Fig. 1a). This was confirmed by quantitative western blotting experiments where PRDX6 levels were compared for cells with and without treatment with dGAE (Fig. 4a).

**Figure 4.**
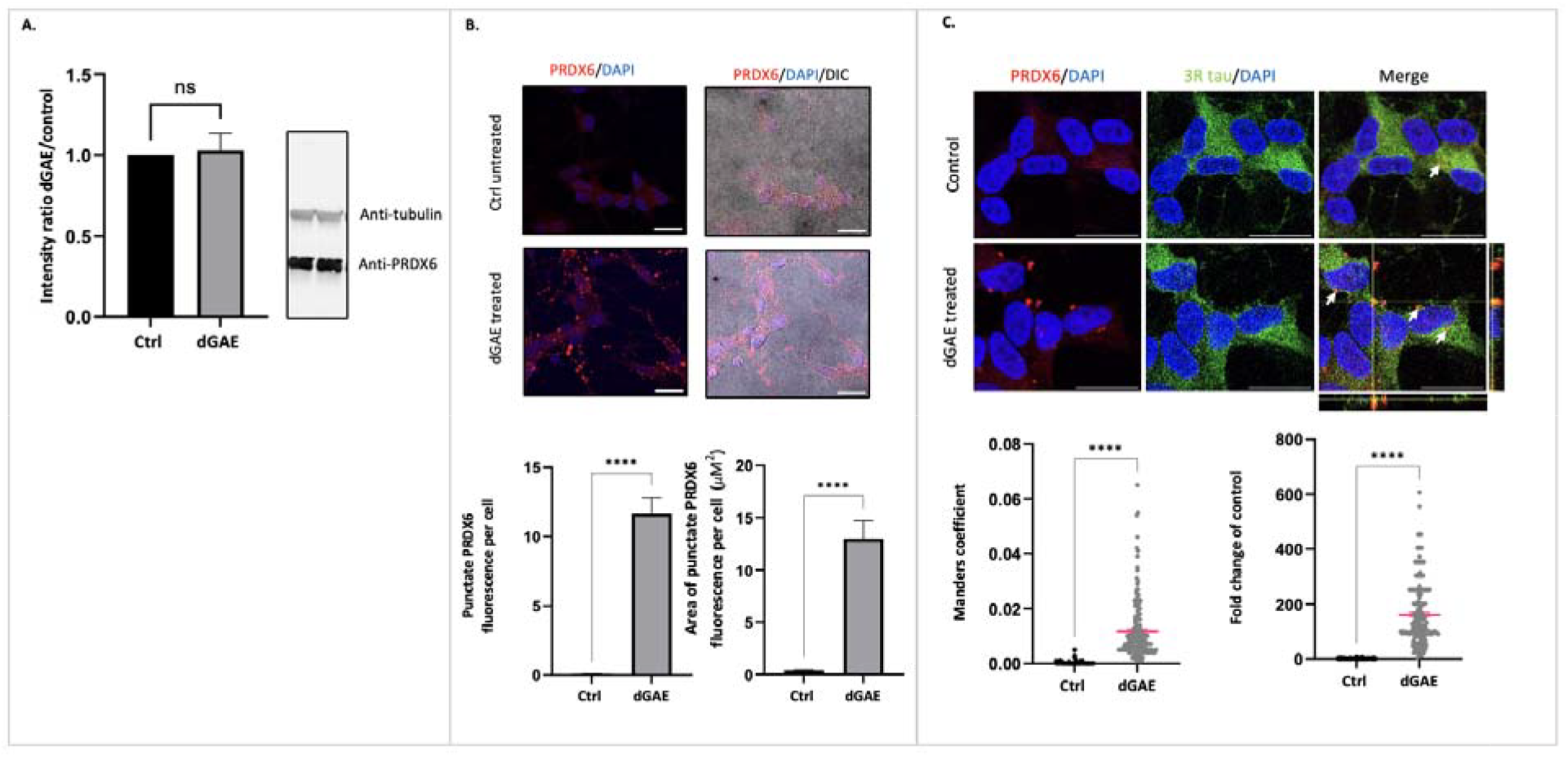
PRDX6 responds to cellular internalisation of dGAE. A) PRDX6 total protein levels are unchanged after internalisation of dGAE. dSH cells were incubated with 10µM dGAE, alongside untreated control cells. Cells were lysed with either RIPA buffer or 4% SDS, separated by SDS-PAGE and visualised by immunoblotting with antibodies to PRDX6 and tubulin. Levels of PRDX6 expression were quantified relative to tubulin and plotted using GraphPad Prism (n=3). B) Images showing dSH cells immunolabelled for PRDX6 (red) with and without treatment of 10µM soluble dGAE for 24h. The Z-stack has been compiled using the Z-project function in FIJI with maximum intensity projection type. All scale bars represent 20µm. Quantification of the particle count of punctate PRDX6 fluorescence normalised to per cell. There were significant differences between the particle count of PRDX6 punctate fluorescence between dGAE (12.98 ± 1.737) and untreated control (0.4047 ± 0.07318) (Mann-Whitney; p < 0.0001). Quantification of the area of punctate PRDX6 fluorescence per cell showed a significant difference between the area of PRDX6 punctate fluorescence between dGAE (11.65µM^2^ ± 1.143µM^2^) and untreated control (0.08331µM^2^ ± 0.01964µM^2^) (Mann-Whitney; p < 0.0001). Confocal analysis is from 7 independent experiments, control = 1435 cells and dGAE = 1964 cells. C) Images showing dSH cells immunolabelled for PRDX6 (red) and endogenous tau with anti-3R tau antibody (green), with and without treatment of 10µM soluble dGAE for 24h. One Z-slice from the centre of the cell is shown. Orthogonal view is displayed for the dGAE treated group to show potential colocalization of endogenous tau and PRDX6. The Z-stacks of images from the control and treated samples were processed and analysed using the BIOP JACoP plugin in FIJI to investigate the co-localisation of PRDX6 and 3R tau fluorescence between the two groups. All scale bars represent 20µm. Colocalization of endogenous tau and PRDX6 was confirmed using the BIOP JACoP in FIJI to measure the Mander’s correlation coefficient between the red and green channels (n= 1090 cells for control, 1149 cells for dGAE from 4 independent tests) There was a significant increase in colocalization between endogenous tau (anti-3R) and PRDX6 between dGAE treated cells (0.01176 ± 0.0008230) and untreated the control (0.0002044 ± 0.00005295) (Mann-Whitney; p < 0.0001). Pink line represents mean with SEM bars. Colocalization after dGAE treatment as a fold-change of the control showing a significant increase in colocalization as a fold-change of control. dGAE treated sample (161.0 ± 8.930) compared to control (1.00 ± 0.1260). Pink line represents mean with SEM bars.

Immunofluorescence was used to detect changes in the localisation of PRDX6 and its co-localisation with tau protein. Comparison of images showing PRDX6 immunostaining of fixed dSH cells showed an increase in PRDX6 signal 24h after dGAE treatment compared to untreated control cells (Fig. 4b). Control cells showed a weak and diffuse PRDX6 signal which became brighter and more punctate after treatment. Changes in PRDX6 punctate fluorescence were quantified. dGAE treatment results in a significant increase of both punctate PRDX6 fluorescence (Fig. 4c) and the area of punctate PRDX6 fluorescence per cell when compared against the untreated control (Fig. 4d). The PRDX6 response was significantly less when dSH cells were incubated with a cysteine mutant of dGAE (dGAE-C322A) (Sup. 7). This suggests that the PRDX6 response requires the presence of dGAE C322 and is not a general response to the internalisation of any peptide.

Co-localisation between endogenous tau and PRDX6 after treatment with dGAE was further detailed by immunolabelling both PRDX6 and endogenous tau. The latter was labelled using anti-3R tau antibody to label amino acids 209-224 of human tau, which recognises endogenous tau in dSH cells but whose epitope is absent from dGAE. The images show co-localization of the red and green channels in the dGAE treated cells (fig. 4e). There was a robust PRDX6 response in the dGAE treated group that shows areas of co-localization of dGAE and PRDX6 labelled with the white arrows. This co-localization was measured using Manders coefficient readout with the BIOP JACoP plugin in Fiji, which illustrates a significant increase in co-localization of the 3R-tau fluorescence with PRDX6 fluorescence after internalisation of dGAE when compared to the control (Fig. 4f). When this data is represented as a fold change of the untreated control (Fig. 4g), there was a 150-fold-change, similar to the observation in proteomic interaction data (Sup. 3). These data support the observation that internalisation of dGAE by dSH cells increases the interaction between PRDX6 and endogenous tau.

### Tau binds to chromatin in dSH cells concurrent with an increase in reactive oxygen species

Tau has been previously shown to associated with chromatin in response to increased OS [3] [22] [23]. Here, we observe both an increase in chromatin proteins in tau immunoprecipitates (Figure 2.a, b) and an increase in co-localization between tau and the antioxidant PRDX6 [24] (Figure 4). To investigate whether increased ROS be important for this change in PRDX6 localisation, we measured changes in ROS levels in dSH cells after internalisation of dGAE using the CellROX™ Green Reagent to detect oxidative stress. dSH cells were treated with 10µM of soluble dGAE for 2h and 24h before being treated with the CellROX™ Green Reagent and fixed for acquisition and quantification in a PerkinElmer Operetta high content imager.

Images of the untreated control cells and dGAE-incubated cells showed similar green fluorescence, indicating ROS (figure 5a), whereas quantification of the intensity of nuclear green fluorescence in treated cells showed a subtle, but significant increase in fluorescence with time, when normalised to the untreated control (figure 5b and c). Staurosporine was included as a positive control (Figure S5). This suggests that soluble dGAE induces a significant increase in ROS in dSH cells. The 24h timepoint shows a greater increase in fluorescence when compared to the 2h timepoint, suggesting that ROS production increased with incubation time.

**Figure 5.**
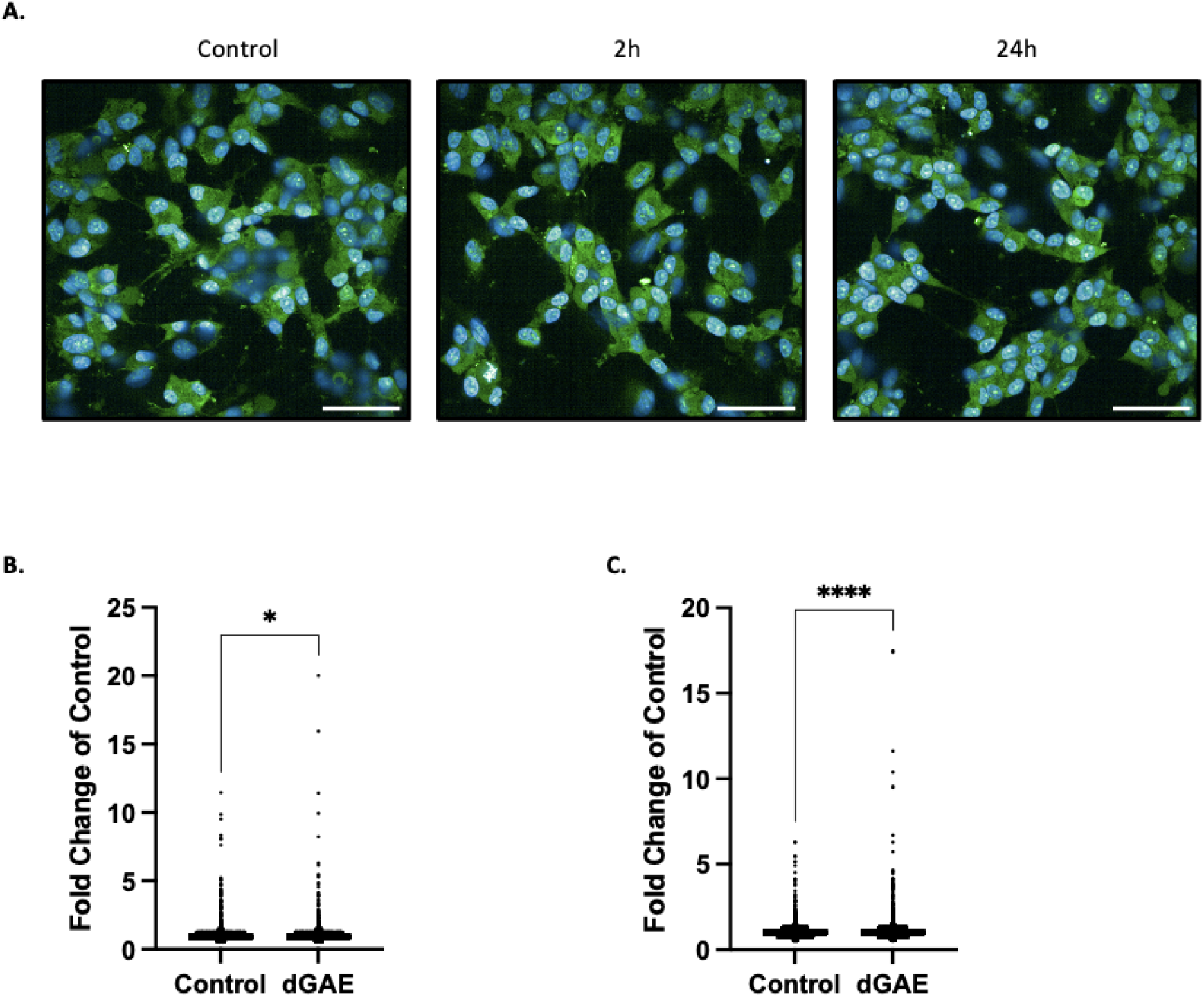
Increased oxidative stress following incubation with soluble dGAE. Untreated dSH cells or cells treated with 10µM soluble dGAE for 2h or 24h were imaged using the CellROX™ Green Reagent to measure oxidative stress. DAPI was used to stain the nucleus (blue) and CellROX™ Green Reagent exhibits bright green photostable fluorescence upon oxidation by reactive oxygen species (ROS) indicating oxidative stress. Scale bar represents 50µm. B) Comparative quantification of CellROX™ Green Reagent after 2h of dGAE treatment. There was a significant difference between the dGAE (2h treatment) (1.008 ± 0.0012) and untreated control (1.003 ± 0.0011) (Mann-Whitney; p = 0.0115). Analysis is from 3 independent experiments, control = 36854 cells and dGAE = 35983 cells. C) Comparative quantification of CellROX™ Green Reagent within after 24h of dGAE treatment. There was a significant difference between the dGAE (24h treatment) (1.021 ± 0.0012) and untreated control (1.003 ± 0.0009) (Mann-Whitney; p < 0.0001). Analysis is from 3 independent experiments, control = 36256 cells and dGAE = 41884 cells.

## Discussion

We have used a proteomic approach to capture global protein alterations in a human neuroblastoma cell line that occur in response to treatment with an amyloidogenic fragment of tau protein, dGAE. It has previously been demonstrated that soluble forms of dGAE are internalised into dSH cells and associate with endogenous tau, leading to its increased phosphorylation and aggregation of the tau [13]. Using the same model system for investigating the propagation of tau pathology, we sought to identify the key pathways or mechanisms that might be involved in early stages of endogenous tau misfolding and modification, utilising the peptide that overlaps the core PHF region [12] [11] as a model system. Interestingly, the mass-spectrometry analysis identified an increase in endogenous tau in response to dGAE treatment which was confirmed using immunoblotting. In our previous study, we showed that internalisation of dGAE led to an increase in insoluble aggregated tau and hyperphosphorylated endogenous tau [13].

To further identify cellular responses to dGAE treatment, interactome analysis was used to identify alterations in endogenous tau binding partners revealing key proteins associated with chromatin, spliceosome and nucleoli indicating that administration of dGAE leads to changes in RNA metabolism and gene expression. There is a growing body of evidence linking tau protein and RNA metabolism and gene expression, with increasing interest in how changes in RNA metabolism might drive neurological pathogenesis of distinct neurological diseases [25] including AD. Tau related changes to gene expression in early Alzheimer’s⍰have been characterized extensively [26] [27] [28]. Tau immunoprecipitation from human brains from either control or AD inputs showed numerous ribonucleoproteins with roles in mRNA processing including splicing and or translation [29]. The authors of the latter study hypothesised that, in AD, the interaction between soluble forms of tau and components of the spliceosome may drive the formation of neurofibrillary tangles. Several heterogeneous nuclear ribonucleoproteins (hnRNP) proteins including HNRNPC have been shown to be upregulated in HEK cells after wild-type tau expression [23]. These mRNA processing proteins are involved in both transcription, translation and alternative splicing, and their dysfunction has been shown to result in cell cycle misregulation with neurological implications leading to neuronal cell death in AD brains [25]. Our data supports a connection between tau and the hnRNP protein HNRNPC. HNRNPC has been shown to both promote the translation of and stabilise the precursor mRNA of the APP gene leading to an increase in Aβ secretion [30] [31].

The abundance of histone proteins in the tau interactome following internalisation of dGAE indicates re-localisation of a subset of tau to the chromatin. Numerous studies have identified a role for tau protein in maintaining chromatin function and genome integrity and to protect cells from DNA damage caused by oxidative stress [3, 22, 32, 33]. Widespread loss of heterochromatin was observed in both tau-transgenic *Drosophila* and mice expressing mutant forms of tau as well as in human AD brains [34]. DNA damage resulting from oxidative stress was identified as the link between transgenic tau expression and the loss of heterochromatin [34]. Additional studies have shown a role for tau in binding to and maintaining chromatin stability [35] [36]. Both acute oxidative stress and heat stress increase the localisation of tau to the nucleus of neuronal cells where it fully protected the neuronal cells against HS-induced DNA damage [3]. In neuronal and neuroblastoma cells, the DNA damaging drug Etoposide increased the translocation of tau to the nucleus suggesting a DNA protective role for tau [37]. In normal mice, cells are protected from the accumulation of DNA oxidative damage after heat shock but tau knock-out mice are not so protected [22]. RNA integrity was also altered after HS conditions in the knock-out mice. A study aimed at understanding the nuclear impact of tau at the transcriptional level, utilising an inducible wild-type or aggregation-prone tau in HEK cells, showed that tau significantly down-regulated several genes implicated in chromatin remodelling and nucleosome organisation during early stages of pathogenesis [23]. Our findings here demonstrating an increase in tau interaction with the chromatin following dGAE treatment may suggest that the binding of tau with chromatin is to protect the DNA from dGAE-induced oxidative stress.

Here we show that nine proteins categorised as nucleolar were enriched in the tau interactome after treatment with dGAE including the rRNA 2’-O-methyltransferase fibrillarin (FBL), a subunit of the first stable ribosome assembly intermediate of the small subunit processome (SSU). The SSU, which is formed inside the nucleolus, is required for RNA folding, modifications, rearrangements, cleavage, and targeted degradation of pre-ribosomal RNA by the RNA exosome [38]. This enrichment of nucleoli proteins in the tau interactome after dGAE internalisation supports a role for tau in ribosomal RNA metabolism and substantiates previous findings [5, 6]. Tau, traditionally recognised for its role in stabilising microtubules, is now known to regulate nuclear and nucleolar processes [33]. We have previously shown that tau localises to the nucleolus in dSH, where it interacts with TIP5, a key regulator of heterochromatin stability and rDNA transcription repression [6]. The enrichment of nucleolar proteins in the tau interactome following dGAE treatment suggests that stress-induced changes may potentially impact the role for tau in rDNA transcription or rRNA processing. This connection is also consistent with our previous observations on the impact of Aβ42 oligomers in dSH where we found that Aβ-induced oxidative stress led to nucleolar tau redistribution associated with alteration in rDNA transcription, rRNA processing and changes in nucleolar proteins such as FBL [5]. Similarly, dGAE treatment appears to induce changes in nucleolar composition, as evidenced by the presence of nucleolar proteins in the tau interactome, suggesting a comparable response to cellular stress. Nucleolar dysfunction in AD, has been widely reported, occurring early before full-blown neurodegeneration, with several nucleolar proteins being dysregulated [39], [40] [41] [42].

Finally, we report increased levels of both PRDX6 and the DNA damage repair protein Parp1 copurifying with tau within 24 hours of treatment of dSH cells with dGAE. This suggests an early role for these proteins in this cellular model for AD, possibly responding to OS signals within the cell. Parp1 is activated by both oxidative stress and DNA damage and has been implicated in the development of neurodegeneration and ageing [43]. The activation of Parp1 can promote both Aβ self-assembly and the formation of tau tangles contributing to the pathogenesis of tau tangles. This protein co-localises with tau, Aβ and microtubule-associated 2 (MAP2) in the brains of AD patients [44]. Here we highlight an interaction between tau, Parp1 and several other proteins including the antioxidant protein PRDX6 and the PRDX6 regulator protein NMP1. Multiple studies have shown an interaction between Parp1 and PRDX6 in various disease states. Several studies have shown altered levels of peroxiredoxin (PRDX) proteins in the brains of patients with distinct neurodegenerative diseases including AD, Parkinson’s disease, Picks disease and Huntington’s disease. [45] [46]. Expression of the antioxidant enzyme PRDX6 has been detected in human astrocytes and at lower levels in neurons where it responds to increases in ROS, resulting from oxidative stress [45]. In this study, we observed a significant increase in PDRX6 fluorescence following dGAE incubation as well as a mild increase in OS.

Since OS has been shown to play a key role in tauopathies, the use of antioxidants might be beneficial for the treatment of tau-related neurodegenerative diseases. This would require an understanding of the molecular mechanisms underlying OS in tauopathies. Proteomic studies are a powerful tool for identifying key changes in pathways or mechanisms that might occur at different stages of neurodegenerative disease. As we have shown here, applying a similar approach to an in vitro model system to study the spreading of tau pathology can provide valuable insight into the early changes that might be taking place as amyloidogenic proteins misfold, revealing targets for therapeutic intervention to prevent the onset or progression of disease.

## References

1. Goedert, M., et al., Multiple isoforms of human microtubule-associated protein tau: sequences and localization in neurofibrillary tangles of Alzheimer’s disease. Neuron, 1989. 3(4): p. 519–26.

2. Crowther, T., M. Goedert, and C.M. Wischik, The repeat region of microtubule-associated protein tau forms part of the core of the paired helical filament of Alzheimer’s disease. Ann Med, 1989. 21(2): p. 127–32.

3. Sultan, A., et al., Nuclear tau, a key player in neuronal DNA protection. J Biol Chem, 2011. 286(6): p. 4566–75.

4. da Costa, P.J., et al., Tau mRNA Metabolism in Neurodegenerative Diseases: A Tangle Journey. Biomedicines, 2022. 10(2).

5. Maina, M.B., et al., The Involvement of Abeta42 and Tau in Nucleolar and Protein Synthesis Machinery Dysfunction. Front Cell Neurosci, 2018. 12: p. 220.

6. Maina, M.B., et al., The involvement of tau in nucleolar transcription and the stress response. Acta Neuropathol Commun, 2018. 6(1): p. 70.

7. Sbodio, J.I., S.H. Snyder, and B.D. Paul, Redox Mechanisms in Neurodegeneration: From Disease Outcomes to Therapeutic Opportunities. Antioxid Redox Signal, 2019. 30(11): p. 1450–1499.

8. Bartolome, F., E. Carro, and C. Alquezar, Oxidative Stress in Tauopathies: From Cause to Therapy. Antioxidants (Basel), 2022. 11(8).

9. Pevalova, M., et al., Post-translational modifications of tau protein. Bratisl Lek Listy, 2006. 107(9-10): p. 346–53.

10. Al-Hilaly, Y.K., et al., Alzheimer’s Disease-like Paired Helical Filament Assembly from Truncated Tau Protein Is Independent of Disulfide Crosslinking. J Mol Biol, 2017. 429(23): p. 3650–3665.

11. Lovestam, S., et al., Assembly of recombinant tau into filaments identical to those of Alzheimer’s disease and chronic traumatic encephalopathy. Elife, 2022. 11.

12. Wischik, C.M., et al., Isolation of a fragment of tau derived from the core of the paired helical filament of Alzheimer disease. Proc Natl Acad Sci U S A, 1988. 85(12): p. 4506–10.

13. Pollack, S.J., et al., Paired Helical Filament-Forming Region of Tau (297-391) Influences Endogenous Tau Protein and Accumulates in Acidic Compartments in Human Neuronal Cells. J Mol Biol, 2020. 432(17): p. 4891–4907.

14. Herrmann, C., D.C. Avgousti, and M.D. Weitzman, Differential Salt Fractionation of Nuclei to Analyze Chromatin-associated Proteins from Cultured Mammalian Cells. Bio Protoc, 2017. 7(6).

15. Luo, Y., et al., Thiosulfate sulfurtransferase deficiency promotes oxidative distress and aberrant NRF2 function in the brain. Redox Biol, 2023. 68: p. 102965.

16. Miaczynska, M., et al., APPL proteins link Rab5 to nuclear signal transduction via an endosomal compartment. Cell, 2004. 116(3): p. 445–56.

17. Zhang, X., et al., Role of Rab GTPases in Alzheimer’s Disease. ACS Chem Neurosci, 2019. 10(2): p. 828–838.

18. Bidinosti, M., et al., Postnatal deamidation of 4E-BP2 in brain enhances its association with raptor and alters kinetics of excitatory synaptic transmission. Mol Cell, 2010. 37(6): p. 797–808.

19. Li, X., et al., Phosphorylated eukaryotic translation factor 4E is elevated in Alzheimer brain. Neuroreport, 2004. 15(14): p. 2237–40.

20. Szklarczyk, D., et al., STRING v10: protein-protein interaction networks, integrated over the tree of life. Nucleic Acids Res, 2015. 43(Database issue): p. D447–52.

21. Chong, Z.Z., F. Li, and K. Maiese, Stress in the brain: novel cellular mechanisms of injury linked to Alzheimer’s disease. Brain Res Brain Res Rev, 2005. 49(1): p. 1–21.

22. Violet, M., et al., A major role for Tau in neuronal DNA and RNA protection in vivo under physiological and hyperthermic conditions. Front Cell Neurosci, 2014. 8: p. 84.

23. Montalbano, M., et al., Tau Modulates mRNA Transcription, Alternative Polyadenylation Profiles of hnRNPs, Chromatin Remodeling and Spliceosome Complexes. Front Mol Neurosci, 2021. 14: p. 742790.

24. Chhunchha, B., E. Kubo, and D.P. Singh, Switching of Redox Signaling by Prdx6 Expression Decides Cellular Fate by Hormetic Phenomena Involving Nrf2 and Reactive Oxygen Species. Cells, 2022. 11(8).

25. Low, Y.H., et al., Heterogeneous Nuclear Ribonucleoproteins: Implications in Neurological Diseases. Mol Neurobiol, 2021. 58(2): p. 631–646.

26. Gunawardana, C.G., et al., The Human Tau Interactome: Binding to the Ribonucleoproteome, and Impaired Binding of the Proline-to-Leucine Mutant at Position 301 (P301L) to Chaperones and the Proteasome. Mol Cell Proteomics, 2015. 14(11): p. 3000–14.

27. Zhang, Q., et al., Integrated proteomics and network analysis identifies protein hubs and network alterations in Alzheimer’s disease. Acta Neuropathol Commun, 2018. 6(1): p. 19.

28. Kavanagh, T., A. Halder, and E. Drummond, Tau interactome and RNA binding proteins in neurodegenerative diseases. Mol Neurodegener, 2022. 17(1): p. 66.

29. Hsieh, Y.C., et al., Tau-Mediated Disruption of the Spliceosome Triggers Cryptic RNA Splicing and Neurodegeneration in Alzheimer’s Disease. Cell Rep, 2019. 29(2): p. 301–316 e10.

30. Lee, E.K., et al., hnRNP C promotes APP translation by competing with FMRP for APP mRNA recruitment to P bodies. Nat Struct Mol Biol, 2010. 17(6): p. 732–9.

31. Rajagopalan, L.E., et al., hnRNP C increases amyloid precursor protein (APP) production by stabilizing APP mRNA. Nucleic Acids Res, 1998. 26(14): p. 3418–23.

32. Wei, Y., et al., Binding to the minor groove of the double-strand, tau protein prevents DNA from damage by peroxidation. PLoS One, 2008. 3(7): p. e2600.

33. Bukar Maina, M., Y.K. Al-Hilaly, and L.C. Serpell, Nuclear Tau and Its Potential Role in Alzheimer’s Disease. Biomolecules, 2016. 6(1): p. 9.

34. Frost, B., et al., Tau promotes neurodegeneration through global chromatin relaxation. Nat Neurosci, 2014. 17(3): p. 357–66.

35. Klein, H.U., et al., Epigenome-wide study uncovers large-scale changes in histone acetylation driven by tau pathology in aging and Alzheimer’s human brains. Nat Neurosci, 2019. 22(1): p. 37–46.

36. Rico, T., et al., Tau Stabilizes Chromatin Compaction. Front Cell Dev Biol, 2021. 9: p. 740550.

37. Ulrich, G., et al., Phosphorylation of nuclear Tau is modulated by distinct cellular pathways. Sci Rep, 2018. 8(1): p. 17702.

38. Sharma, D., D.D. Licatalosi, and E. Jankowsky, Kinetics of RNA-protein interactions in cells. Trends Biochem Sci, 2021. 46(10): p. 861–862.

39. Nyhus, C., et al., Evidence for nucleolar dysfunction in Alzheimer’s disease. Rev Neurosci, 2019. 30(7): p. 685–700.

40. Garcia-Esparcia, P., et al., Altered machinery of protein synthesis is region- and stage-dependent and is associated with alpha-synuclein oligomers in Parkinson’s disease. Acta Neuropathol Commun, 2015. 3: p. 76.

41. Ding, Q., et al., Ribosome dysfunction is an early event in Alzheimer’s disease. J Neurosci, 2005. 25(40): p. 9171–5.

42. Ding, Q., et al., Decreased RNA, and increased RNA oxidation, in ribosomes from early Alzheimer’s disease. Neurochem Res, 2006. 31(5): p. 705–10.

43. Mao, K. and G. Zhang, The role of PARP1 in neurodegenerative diseases and aging. FEBS J, 2022. 289(8): p. 2013–2024.

44. Narne, P., et al., Poly(ADP-ribose)polymerase-1 hyperactivation in neurodegenerative diseases: The death knell tolls for neurons. Semin Cell Dev Biol, 2017. 63: p. 154–166.

45. Krapfenbauer, K., et al., Aberrant expression of peroxiredoxin subtypes in neurodegenerative disorders. Brain Res, 2003. 967(1-2): p. 152–60.

46. Szeliga, M., Peroxiredoxins in Neurodegenerative Diseases. Antioxidants (Basel), 2020. 9(12).

