## Supplementary data for "A proteomic analysis of the PHF-forming tau fragment (tau297-391) following uptake into differentiated human neuronal SHSY5Y cells"

**Supplementary Figures**

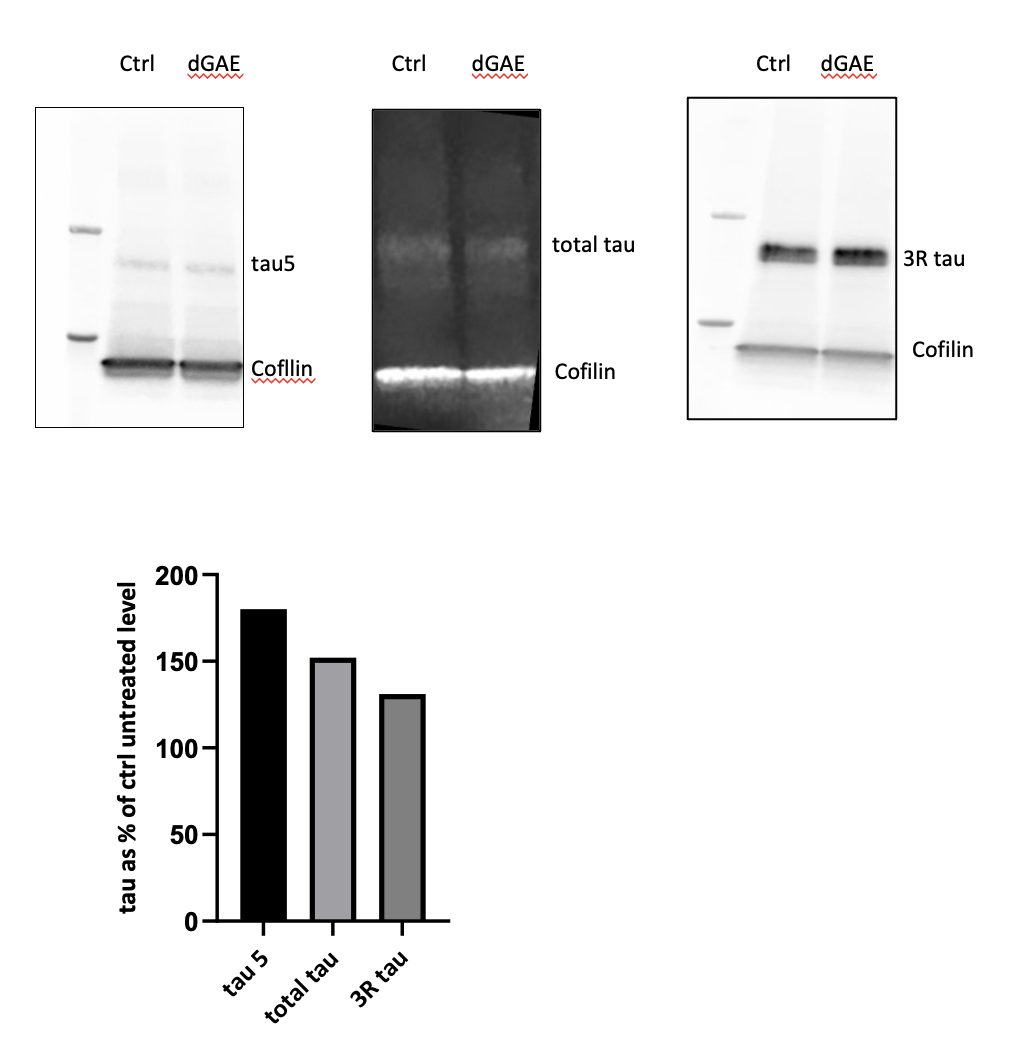

***Supplementary Figure 1.*** ***Tau expression levels.*** *Protein lysates were prepared from dSH cells 24 hours post incubation with 10µM dGAE, alongside untreated control cells. Protein was lysed using RIPA buffer, separated by SDS-PAGE and visualised by immunoblotting with antibodies to tau (MAPT) and cofilin.* *Levels of tau expression were quantified relative to cofilin loading control and plotted using GraphPad Prism.*

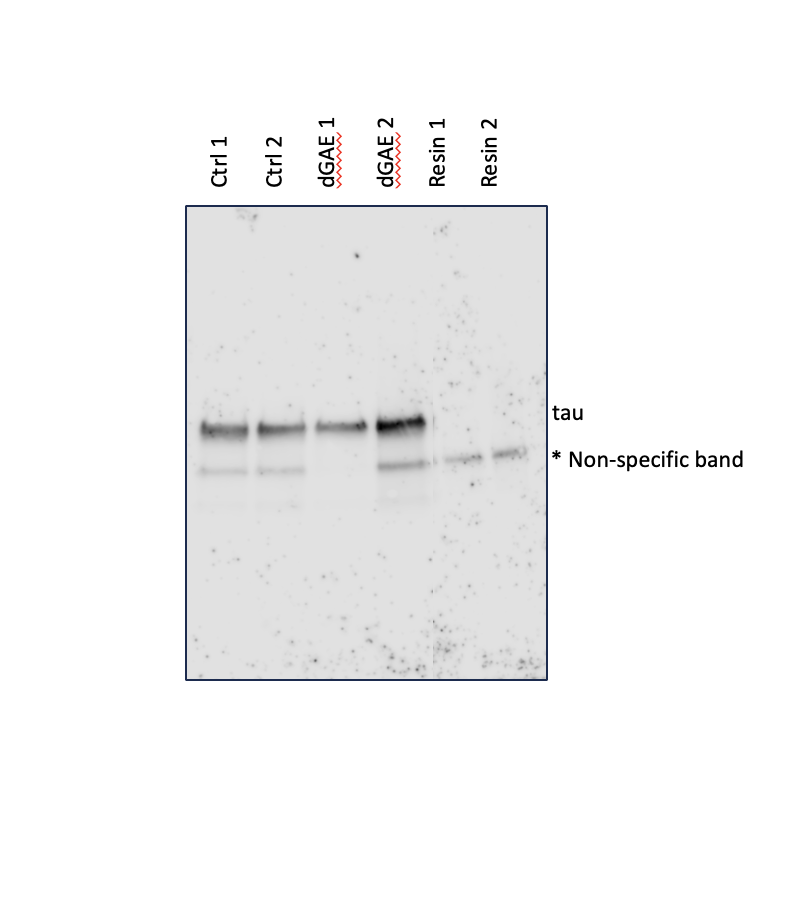

***Supplementary*** ***Figure 2.*** ***Tau co-immunoprecipitation****. Tau and co-precipitating proteins were purified from whole cell lysates using antibodies to total tau (Rabbit, SAB4501831, Sigma) as described in the materials and method section. Eluted protein from control untreated, 10µM dGAE treated (24 hours) and resin only samples was separated by SDS-PAGE and visualised by immunoblotting with antibodies to tau (Mouse MAB3420, Chemicon).*

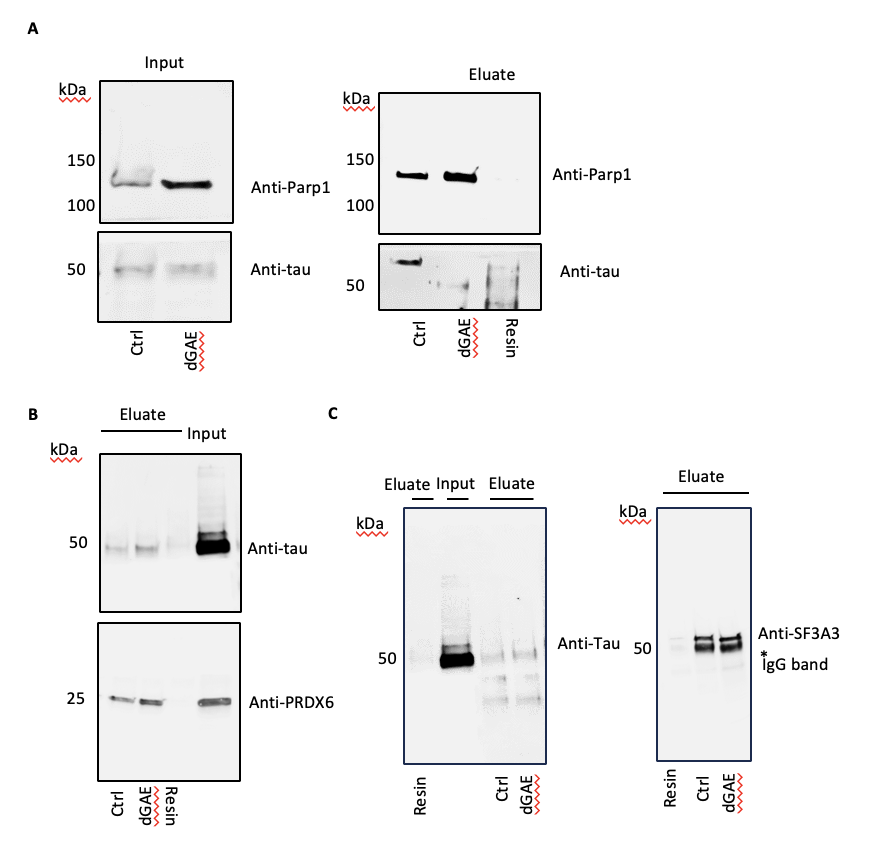

***Supplementary Figure 3. Tau co-purifies with parp1, SF3A3 and PRDX6****.* *Protein lysates were prepared for immunoprecipitation from dSH cells 24 hours post incubation with 10µM dGAE, alongside untreated control cells. Cells were lysed in IP buffer and incubated with either parp1 antibody (46D11 cell Signalling), PRDX6 antibody (Proteintech 13585-1-AP) or SF3A3 antibody* *(proteintech 12070-1-AP) as described in the materials and methods section. Precipitates from control untreated, dGAE treated and resin controls were eluted into 7M Urea, 2M Thio-urea, separated by SDS-PAGE and visualised by western blotting using antibodies to parp1 (46D11 cell Signalling) PRDX6 (Proteintech 13585-1-AP), SF3A3 (proteintech 12070-1-AP) and tau (Mouse, MAB3420, Chemicon).* *Input samples are shown for antibody controls.* ***A.*** *Parp1 immunoprecipitation.* ***B.*** *PRDX6 immunoprecipitation.* ***C.*** *SF3A3 immunoprecipitates.*

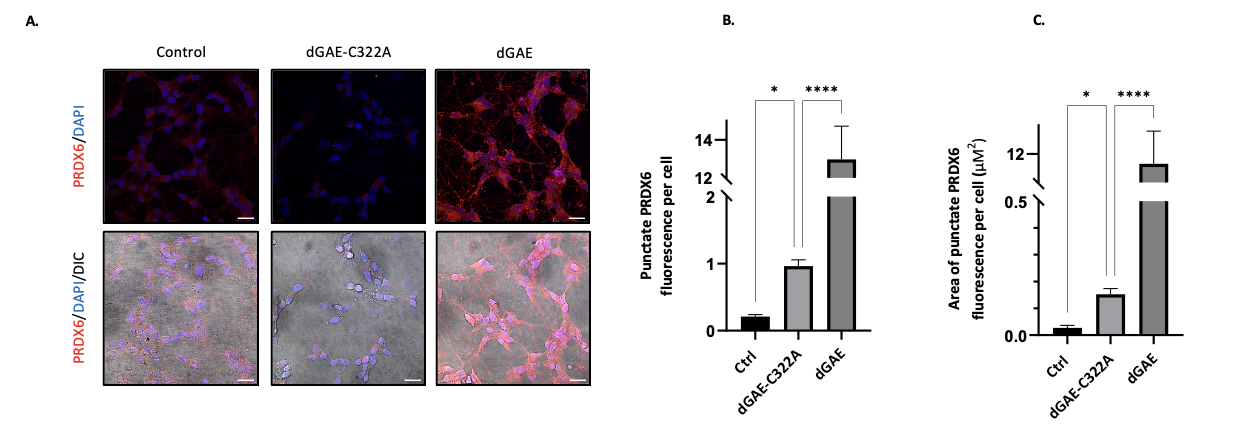

***Supplementary figure 4. PRDX6 response is not observed with the treatment of dGAE-C322A.*** *Confocal images of dSH cells immunolabelled for PRDX6 (red) without treatment and the treatment of 10µM dGAE-C322A or 10µM dGAE for 24h. The Z-stack has been compiled using the Z-project function in FIJI with maximum intensity projection type. (b) Quantification and comparison of the particle count of punctate PRDX6 fluorescence per cells in control, dGAE-C322A and dGAE treated cells. Kruskal-Wallis ANOVA shows there is a significant difference between the groups (p < 0.0001). Dunn’s multiple comparison test shows a significant difference between dGAE-C322A treated (0.9637 ± 0.09505) when compared to the control (0.2112 ± 0.02975, p = 0.0317) and dGAE treated (12.98 ± 1.737, p < 0.0001). (c) Quantification and comparison of the area of punctate PRDX6 fluorescence per cell for control, dGAE-C322A and dGAE treated cells. Kruskal-Wallis ANOVA shows there is a significant difference between the groups (p < 0.0001). Dunn’s multiple comparison test shows a significant difference in dGAE-C322A treated (0.1523μM^2^ ± 0.02162μM^2^) when compared to the control (0.02699μM^2^ ± 0.009585μM^2^, p = 0.0467) and dGAE sample (11.65μM^2^ ± 1.143μM^2^, p < 0.0001). Confocal analysis comes from 4 independent tests for control (1162 cells) and dGAE-C322A (982 cells), 6 independent tests for dGAE (1964 cells).*

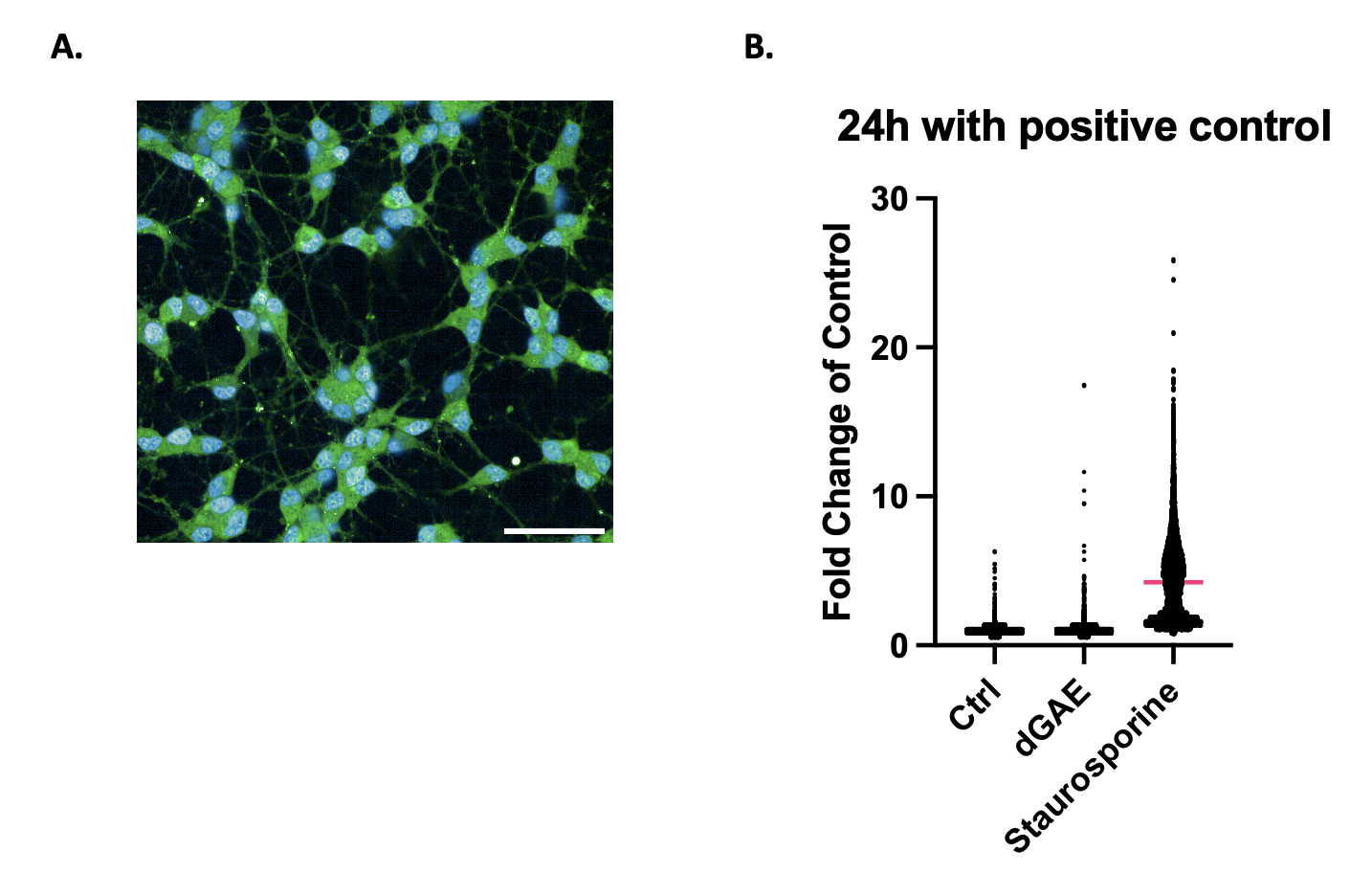

***Supplementary Figure 5****. Positive control to show the effect of staurosporine. A) Image of dSH cells treated with 7µM staurosporine for 1h prior to CellROX™ Green Reagent treatment.*

*B) Quantification of 24h treatment with positive control. Untreated control (1.003 ± 0.0009), dGAE (1.021 ± 0.0012) and staurosporine (4.244 ± 0.025) Analysis is from 3 independent experiments, control = 36256 cells, dGAE = 41884 cells and staurosporine = 13462.*

**Supplementary Tables**

| **Majority protein IDs** | **Protein names** | **Gene names** | **Fold Change_dGAE high** | **Student's T-test p-value Control_dGAE** | **Abbreviated Function from Uniprot** |
| --- | --- | --- | --- | --- | --- |
| P30041 | Peroxiredoxin-6 | PRDX6 | 353.34 | 0.02442 | Cell Protection against oxidative stress |
| Q9ULV4 | Coronin-1C | CORO1C | 13.99 | 0.01803 | Cell Migration |
| P50897 | Palmitoyl-protein thioesterase 1 | PPT1 | 10.92 | 0.00840 | Lysosomal degradation |
| Q12874 | Splicing factor 3A subunit 3 | SF3A3 | 9.60 | 0.04985 | pre-mRNA splicing |
| Q9UKV3 | Apoptotic chromatin condensation inducer in the nucleus | ACIN1 | 7.86 | 0.01556 | Splicing process |
| Q13813 | Spectrin alpha chain, non-erythrocytic 1 | SPTAN1 | 7.50 | 0.02676 | Interacts with Calmodulin |
| Q09028 | Histone-binding protein RBBP4 | RBBP4 | 7.46 | 0.02127 | Chromatin metabolism |
| Q04726 | Transducin-like enhancer protein 3 | TLE3 | 7.17 | 0.03176 | Transcriptional corepressor |
| P31942 | Heterogeneous nuclear ribonucleoprotein H3 | HNRNPH3 | 6.88 | 0.00497 | Splicing process |
| Q02809 | Procollagen-lysine,2-oxoglutarate 5-dioxygenase 1 | PLOD1 | 6.11 | 0.04779 | Assembly and cross-linking of Collagen fibrils |
| Q7RTV0 | PHD finger-like domain-containing protein 5A | PHF5A | 5.80 | 0.03088 | component of the splicing factor SF3B complex |
| P22087 | rRNA 2-O-methyltransferase fibrillarin | FBL | 5.30 | 0.01619 | RNA and protein methyltransferase |
| Q9Y3Y2 | Chromatin target of PRMT1 protein | CHTOP | 4.96 | 0.00419 | Includes associates with spliced mRNA as part of TREX comlex |
| Q9Y5B9 | FACT complex subunit SPT16 | SUPT16H | 4.92 | 0.02971 | Chromatin factor that acts to reorganise nucleosomes |
| Q15428 | Splicing factor 3A subunit 2 | SF3A2 | 4.89 | 0.04983 | pre-mRNA splicing |
| P09234 | U1 small nuclear ribonucleoprotein C | SNRPC | 4.42 | 0.00961 | pre-mRNA splicing and assembly of Spliceasome |
| P09874 | Poly [ADP-ribose] polymerase 1 | PARP1 | 4.167 | 0.03664 | Poly-ADP-ribosyltransferase which plays a key role in DNA repair |
| Q7Z7K6 | Centromere protein V | CENPV | 3.96 | 0.01279 | Chromatin dynamics |
| P09382 | Galectin-1 | LGALS1 | 3.77 | 0.008737 | Plays a role in regulating apoptosis, cell proliferation and cell differentiation |
| P18124 | 60S ribosomal protein L7 | RPL7 | 3.76 | 0.027987 | Component of large ribosomal subunit |
| P07910 | Heterogeneous nuclear ribonucleoproteins C1/C2 | HNRNPC | 3.63 | 0.01970 | Modulates the translation of bound mRNA molecules |
| Q16778 | Histone H2B type 2-E | HIST2H2BE | 3.60 | 0.03777 | Core component of Nucleosome |
| Q13435 | Splicing factor 3B subunit 2 | SF3B2 | 3.51 | 0.00997 | pre-mRNA splicing |
| P12814 | Alpha-actinin-1 | ACTN1 | 3.13 | 0.02408 | F-actin cross-linking protein |
| Q9BZE4 | Nucleolar GTP-binding protein 1 | GTPBP4 | 3.04 | 0.02145 | Involved in the biogenesis of the 60S ribosomal subunit and role in cell cylcle arrest |
| Q99878 | Histone H2A type 1-J | HIST1H2AJ | 2.95 | 0.03521 | Core component of Nucleosome |
| P46821 | Microtubule-associated protein 1B | MAP1B | 2.91 | 0.04634 | microtubule dynamics and role in neuronal differentiation and neurite extension |
| Q15393 | Splicing factor 3B subunit 3 | SF3B3 | 2.78 | 0.00533 | pre-mRNA splicing |
| Q14498 | RNA-binding protein 39 | RBM39 | 2.71 | 0.03843 | pre-mRNA splicing |
| Q99879 | Histone H2B type 1-M | HIST1H2BM | 2.62 | 0.01471 | Core component of Nucleosome |
| O00159 | Unconventional myosin-Ic | MYO1C | 2.62 | 0.04661 | Unconventional myosins serve in intracellular movements |
| Q71UI9 | Histone H2A.V | H2AFV | 2.43 | 0.00456 | Variant Histone H2A replaces conventional H2A in a subset of nucleosomes |
| Q9NWH9 | SAFB-like transcription modulator | SLTM | 2.36 | 0.04509 | General inhibitor of transcription that eventually leads to apoptosis |
| P46777 | 60S ribosomal protein L5 | RPL5 | 2.29 | 0.0208 | Component of the ribosome |
| P55769 | NHP2-like protein 1 | NHP2L1 | 2.20 | 0.00566 | pre-mRNA splicing |
| Q9UHB9 | Signal recognition particle subunit SRP68 | SRP68 | 2.18 | 0.01801 | Component of the SRP, targetting of secretory and membrane proteins to ER |
| P06748 | Nucleophosmin | NPM1 | 2.06 | 0.01613 | Involved in diverse cellular processes |
| Q15717 | ELAV-like protein 1 | ELAVL1 | 2.03 | 0.0280 | binds to the 3'-UTR region of mRNAs and increases their stability |
| P40425 | Pre-B-cell leukemia transcription factor 2 | PBX2 | 2.016 | 0.00913 | Transcriptional activator |

**Supplementary Table 1 to show proteins enriched in tau immuno-precipitates after dGAE treatment.** Changes to the tau interactome in dSH cells were explored using co-immunoprecipitation combined with quantitative LC-MS as described in the materials and methods section. T test analysis was used to identify changes to the tau interactome. Proteins that were enriched two-fold or more after dGAE treatment with a P value of <0.05 were selected.

| **#term ID** | **term description** | **observed gene count** | **background gene count** | **strength** | **false discovery rate** |
| --- | --- | --- | --- | --- | --- |
| GO:0070013 | Intracellular organelle lumen | 37 | 5857 | 0.49 | 2.68E-13 |
| GO:0032991 | Protein-containing complex | 33 | 5073 | 0.5 | 5.05E-11 |
| GO:0031981 | Nuclear lumen | 32 | 4733 | 0.52 | 6.27E-11 |
| GO:1990904 | Ribonucleoprotein complex | 15 | 677 | 1.03 | 7.10E-10 |
| GO:0005634 | Nucleus | 35 | 7390 | 0.36 | 2.69E-08 |
| GO:0005654 | Nucleoplasm | 27 | 3973 | 0.52 | 3.24E-08 |
| GO:0005681 | Spliceosomal complex | 9 | 192 | 1.36 | 4.10E-08 |
| GO:0005684 | U2-type spliceosomal complex | 7 | 94 | 1.56 | 2.15E-07 |
| GO:0071005 | U2-type precatalytic spliceosome | 6 | 50 | 1.77 | 2.27E-07 |
| GO:0043232 | Intracellular non-membrane-bounded organelle | 28 | 4880 | 0.45 | 3.60E-07 |
| GO:0097525 | Spliceosomal snRNP complex | 6 | 56 | 1.72 | 3.60E-07 |
| GO:0016604 | Nuclear body | 13 | 789 | 0.91 | 3.74E-07 |
| GO:0043229 | Intracellular organelle | 40 | 12528 | 0.19 | 1.65E-06 |
| GO:0005686 | U2 snRNP | 4 | 18 | 2.04 | 8.99E-06 |
| GO:0005730 | Nucleolus | 12 | 924 | 0.8 | 1.71E-05 |
| GO:0043231 | Intracellular membrane-bounded organelle | 37 | 10761 | 0.23 | 1.94E-05 |
| GO:0043227 | Membrane-bounded organelle | 39 | 12427 | 0.19 | 2.35E-05 |
| GO:0071013 | Catalytic step 2 spliceosome | 5 | 87 | 1.45 | 8.01E-05 |
| GO:0016607 | Nuclear speck | 8 | 399 | 0.99 | 9.04E-05 |
| GO:0032993 | protein-DNA complex | 6 | 195 | 1.18 | 0.00019 |
| GO:0001651 | Dense fibrillar component | 2 | 2 | 2.69 | 0.0014 |
| GO:0005689 | U12-type spliceosomal complex | 3 | 25 | 1.77 | 0.0014 |
| GO:0070062 | Extracellular exosome | 14 | 2099 | 0.51 | 0.0023 |
| GO:0000786 | Nucleosome | 4 | 106 | 1.27 | 0.0037 |
| GO:0031428 | Box C/D snoRNP complex | 2 | 7 | 2.15 | 0.0068 |
| GO:0005925 | Focal adhesion | 6 | 405 | 0.86 | 0.0079 |
| GO:0005694 | Chromosome | 11 | 1712 | 0.5 | 0.0215 |
| GO:0071004 | U2-type prespliceosome | 2 | 19 | 1.71 | 0.0353 |

***Supplementary Table 2. A PPI network of the enriched proteins identified in a T test comparing dGAE treated to untreated control cells.*** *Proteins that were enriched two-fold or more after dGAE treatment with a P value of <0.05 were selected. The PPI network* *was generated for enriched proteins using the STRING database. Functional enrichments for the uploaded proteins are shown for Gene Ontology terms classified as cellular component.*

| **Protein names** | **Gene names** | **Fold Change_Control high** | **Student's T-test p-value dGAE to Control** | **Abbreviated Function from Uniprot** |
| --- | --- | --- | --- | --- |
| Vacuolar protein sorting-associated protein 54 | VPS54 | 1292.463922 | 0.01288001 | Transport from Endosomes to Golgi network |
| Kallikrein-5 | KLK5 | 22.07286009 | 0.037938769 | May be involved in desquamation |
| Statherin | STATH | 19.74561733 | 0.088392394 | inhibits precipitation of calcium phosphate salts |
| Proactivator polypeptide-like 1 | PSAPL1 | 6.914502848 | 0.032962957 | May activate the lysosomal degradation of sphingolipids |
| Y-box-binding protein 2 | YBX2 | 5.215236328 | 0.014792386 | Regulation of the stability and/or translation of germ cell mRNAs |
| Extracellular glycoprotein lacritin | LACRT | 3.787071307 | 0.00563755 | Modulates secretion by lacrimal acinar cells |
| Retroviral-like aspartic protease 1 | ASPRV1 | 3.759237307 | 0.022400433 | Protease responsible for filaggrin processing |
| Ig lambda chain V-III region LOI | IGLV3-9 | 3.700231877 | 0.027379345 | Immunoglobulins |
| Lactotransferrin | LTF | 3.416872202 | 0.041241683 | Iron binding transport protein |
| Lysozyme C | LYZ | 3.386256035 | 0.047736034 | Lysozymes |

**Supplementary Table 3. Proteins enriched in tau immuno-precipitates from untreated control cells compared to dGAE treated cells.** Changes to the tau interactome in dSH cells were explored using co-immunoprecipitation combined with quantitative LC-MS as described in the materials and methods section. T test analysis was used to identify changes to the tau interactome. Proteins that were enriched two-fold or more in control untreated cells compared to dGAE treated cells with a P value of <0.05 were selected.
